# New Protein Function Characterization for Human Paralog Discovery, Scraping the Bottom of the Genomics Barrel

**DOI:** 10.64898/2026.05.28.728578

**Authors:** BK Pradeep, Weixia Deng, Robert L. Jernigan

## Abstract

Increasing the number of related protein paralogs is important for fully understanding protein relationships, yet it remains challenging for sequences in the twilight zone. Here, we present an integrated homolog detection framework that combines sequence-based (BLASTp, MMseqs2), structure-based (Foldseek), and embedding-distance-based (PROST) similarity metrics to identify additional paralogs. To characterize functionally related protein pairs, we develop protein-family-specific supervised logistic regression models trained on curated functional annotations from MEROPS proteases and KinHub kinases. The resulting model successfully classifies proteins, with ROC-AUC of 0.99 and F1-score of 0.92 for test datasets. Applying this model, we initially identify 686 protease and 298 kinase new candidates. Subsequent structural validation, and previous annotation comparisons yield 7 new protease and 3 new kinase paralogs in the human proteome, mostly lacking prior functional characterization. An additional outcome is structural identification of catalytically important residues for larger numbers of proteases and kinases. Despite the small number of new paralogs for well-studied proteases and kinases, our results demonstrate that integrating orthogonal homolog approaches with family-specific regression models provides a robust, scalable strategy for discovering new functionally related proteins, which is a generalizable approach for novel protein function discovery and can be applied more broadly to under-annotated proteomes.

## Introduction

Advances in sequencing technologies have resulted in the unprecedented explosion in the available sequence data[1,2]. Also, huge numbers of genome sequences are now available, including metagenomic sequence that contribute billions of sequence information [3]. Generally, sequence comparisons are used to compare proteins to identify the similarities and differences between them. When pairs are sufficiently similar than functions can be inferred for the target. Identifying homologs in this way is fundamental for understanding evolution, function and mechanism. Because protein structures are more conserved than sequences, structure comparisons provide additional confirmation of homolog relationships. High levels of structure similarity are always found for homologs, even for those within the twilight zone of sequence identity [4,5]. In the present work, homologs will be identified, and their predicted structures will be compared to validate the homolog relationships and the transferred functions from annotated homologs for the novel homologs. Also, breakthroughs in AI-driven deep learning protein structure predictions such as AlphaFold [6,7], ESMFold [8], RoseTTAFold [9], DeepFold [10], OmegaFOld [11], ProteinBERT [12] have made available relatively reliable predicted structures of almost every protein, with accuracies approaching those of experimental methods. The AlphaFold database [6,7] alone hosts over 200 million predicted protein structures. Despite these remarkable advances, functional annotations remain incomplete for a significant fraction of proteins[13]. For example, only about half-a million proteins are manually curated in the Swiss-Prot database[14,15]. Thus, the data available offers an unprecedented opportunity to infer novel functional relationships among proteins at the whole proteome scale.

Traditionally, sequence-based homolog search methods such as BLASTp [16] have been widely used to identify similar proteins. Faster alternatives including UBLAST [17], RAPSearch2 [18], DIAMOND [19] have been developed to address the increasing scale of sequence databases. More recently, MMseqs2 [2] has emerged as a highly sensitive and fast method for large-scale sequence homolog detection. These sequence-based approaches infer functional relatedness primarily from significant alignment scores. However, many remote homologs cannot be reliably detected by sequence similarity alone. As a result, structure-based methods such as Dali [20], TM-align [21], and CE [22] have been used to identify distantly related proteins. Foldseek enables improved speed and sensitivity for identifying structurally similar protein, and thereby homologs [23]. Some methods such as CATH [24] use both sequence and structure information, to characterize protein domains into different families to establish structural relationships.

Recently, new homolog detection methods have appeared that are utilize the Large Protein Language Models (LPLMs) with a remarkable ability to find new homologs, because they have digested huge number of sequences to capture overall the biophysical, biochemical, evolutionary, phylogenetic, structural, and mutational characteristics within their embeddings [25]. For a simple bacteria like *Bacillus subtillis*, our method PROST [5] enabled a nearly complete set of functional annotations; see https://bit.ly/prost-bsubtilis. Protein Language Models (pLMs) trained on protein sequences (e.g. ESM [8,26,27], ProtT5 [28], protBERT [29], ProGen[30], PLMSearch[31]), structures (TM-Vec[32]), or combined sequence-structure (ProstT5[33], ProteinGPT[34]) generate comprehensive embedding representations that can be leveraged for diverse downstream application such as homology identification, protein function predictions, mutation effect predictions, de-novo protein design, mapping protein interactions, and drug discovery. The effectiveness of these embedding-based representations has also been demonstrated in the CAFA5 [35] protein annotation prediction challenge [35]. Nonetheless, these approaches remain associated with several challenges like high computational demand, data bias, limited interpretability, and fine-tuning complexities[36].

Despite sharing similar sequences or structures, some proteins can exhibit functional differences due to their differences in cellular environment, localization, catalytic sites, and dynamics [37–39]. Some protein families retain conserved structures despite their functional differences as seen in the α/β hydrolases [40]. For instance, α/β hydrolase proteins - serine carboxypeptidases and lipases differ functionally, despite sharing similar structural folds and identical catalytic triads, making functional discrimination among structural homologs sometimes challenging[41,42]. One approach to improve reliable function annotations, GoRetriever uses an information retrieval approach to distill information from literature searches to improve protein annotation[43]. ProtSpace integrates pLM embeddings along with integrated annotations across different databases for improved functional characterization of proteins[44].

The aim of this study is to discover novel protein functions in human proteome. In this study, we present an integrated homolog detection framework that combines sequence (BLAST[16], MMseqs2[2]), structure (Foldseek[23]), and embedding distance-based (PROST[5]) similarity scores to identify functionally related paralog pairs in the human proteome. As proteases and kinases are among the most studied enzymes in humans, we use these families for detailed studies. Using this integrated approach, we first perform all-vs-all homolog searches for three datasets: 20,647 human protein sequences, 675 human proteases, and 522 human kinases from the UniProtKB reference proteome database, the MEROPS database, and the KinHub database, respectively. This gives us huge number of paralog pairs backed by sequence-structure-embedding-based similarity signals but does not provide clear thresholds to define functional closeness. To systematically integrate orthogonal similarity signals and identify functionally similar paralogs, here we employ family-specific supervised logistic regression [45] model. The model estimates the probability of protein pairs belonging to the same functional group. For Foldseek homolog search, we use AlphaFold predicted structures. Then, we train a protein-disjoint logistic regression model on curated functionally similar annotation cases (i.e., MEROPS[46] protease clans, Kinhub[47] kinase family) to learn the optimal combination of these different similarity signals. The supervised training learns to differentiate between functionally related and unrelated protein pairs. The model can only be used within the protein family for which it has been trained, but similar regression models could be developed in the same fashion within or across species for different protein families. To ensure similar functions of these enzymes, we investigate the details in the structures of the corresponding active-site residues against the previously known reference enzymes. Upon manual inspection of annotations, motifs, and catalytic residue conservation, we identify seven novel protease candidates and three novel kinase candidates in humans. Most of these novel findings either lack active-site residue identifications or are not sufficiently annotated as proteases or kinases. Our results demonstrate that the family-specific supervised classifier model integrating multiple homolog scores along with detailed structural comparisons provide a principled and reliable way to identify functionally related protein paralogs.

## Results

### Regression Discrimination for Human Protease and Kinase Homologs Predicted from Integrated Methods

The first step of this study is to find homologs within the human proteome using four different methods. We perform method-specific exhaustive all-vs-all homology searches and find homologs for each query protein. For our study, we analyze homologs for the Swiss-Prot reviewed 536 human proteases and 736 human kinases using our integrated homolog search framework that combines sequence, structure, and embedding distance-based homolog methods. After filtering homolog pairs for reciprocal hits and statistical significance, the initial datasets have 1,305 putative proteases and 2,274 putative kinases that were not annotated previously in the reference Swiss-Prot database. The method-specific distributions and overlaps of homolog pairs are shown for proteases and kinases in **Fig. 1A** and **1B**, respectively. The result demonstrates that most of the homolog pairs are found by MMseqs2 and Foldseek. Moreover, each method uniquely finds paralog pairs, each complementing the homology search, highlighting the advantage of integrating multiple similarity signals.

**Figure 1.**
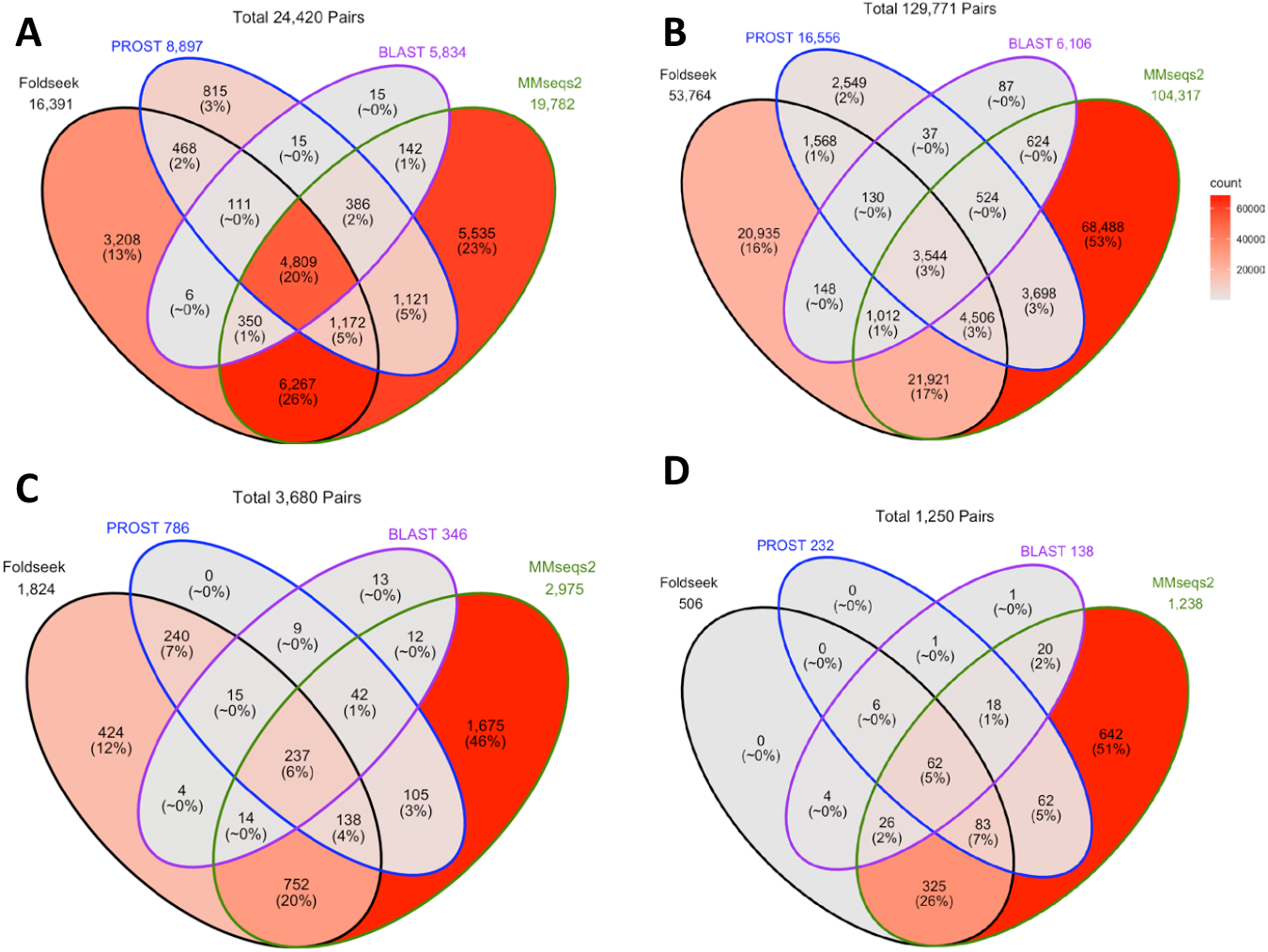
Distribution of novel and validated human protease and human kinase paralog pairs identified by integrated homology searches and regression model. **(A, B)** Initial paralog pairs found by four methods (BLASTp, MMseqs2, Foldseek, and PROST) and those found by multiple methods. (**C, D)** Distribution of putative protease pairs and putative kinase pairs retained after applying protein-family specific regression model trained to predict functional relatedness. A total of 686 novel protease candidates and 298 novel kinase candidates are predicted to be functionally related. Following these, further validation steps are based on catalytic residue conservation.

For further refinement, we apply family-specific trained logistic regression models for the proteases and kinases (**details in Method**) to ensure the functional relatedness among paralog pairs. Only 686 protease candidates and 298 kinase candidates are then predicted to be functionally close. The distributions of these filtered candidate paralogs are shown in **Fig. 1C** and **1D**, respectively. These results indicate that our regression model is conservative and substantially reduces the paralog sets while ensuring candidates have extremely high functional similarity.

### Novel Paralogs with Predicted Catalytic-Residue Conservation

For the functionally related paralog pairs predicted above, we next perform pairwise structural alignments using FoldMason[48] to examine the conservation of mechanistically important active-site residues between the reference proteins and the corresponding novel candidates. From this analysis, we identify 98 candidate proteases and 26 candidate kinases that retain the same catalytic residues as their respective reference proteins.

To further refine our candidate sets, we investigate the conservation of the motifs, particularly for metalloproteases that require preservation of a characteristic HExxH catalytic signature[49–51]. Furthermore, we perform detailed annotation inspection, catalytic-residue mapping, and structural comparisons for these novel candidates. This reduces the sets to seven novel protease candidates and three novel kinase candidates within the human proteome. Interestingly, many of these predicted active site residues are currently not annotated in UniProt and are highlighted in red in **Table 1**. In addition, most of these candidate proteins remain poorly characterized with respect to their potential enzymatic activities, suggesting they may have these functions.

**Table 1.** Active site residue predictions for novel candidates and their validation. Red highlighted are predicted active site residues identified by pairwise structural alignment using Foldmason.

|  |  |  |  |  |  |
| --- | --- | --- | --- | --- | --- |
| **A. Active site residue predictions for novel human proteases against known references** |  |  |  |  |  |
| Reference protein (id) | novel candidate (target id) | active site residues in reference | active-site residues for target | TM-score | Validation for novel candidates |
| Q8TAC2 | Q96NT3 | C24 H125 | **C33 H186** | 0.42 | CATH: in the cysteine proteinases superfamily |
| Q99436 | P28070 | T44 | **T54** | 0.66 | UniProt: non-catalytic component of proteasome complex; conserved local topology around predicted catalytic residues |
| P19440 | P36268 | T381 | T381 | 0.98 | UniProt: Inactive glutathione hydrolase ; high structural and sequence similarity along with conserved active-site residue |
| P13798<br>461-732 | Q5JUR7 | S587<br>D675<br>H707 | **S102 D163**<br>**H192** | 0.66 | InterPro[52]: has $\alpha/\beta$ hydrolase domain<br>UniProt annotation score: 2/5<br>Pfam: high structural alignment with prolyl oligopeptidase domain of reference counterpart with catalytic residue conservation |
| P27487<br>497-766 | Q7L211 | S630<br>D708<br>H740 | S193 D268<br>H298 | 0.70 | UniProt: MF- hydrolase activity<br>InterPro: has $\alpha/\beta$ hydrolase domain<br>Annotation score: 3/5<br>Pfam: structural similarity with prolyl oligopeptidase domain of reference counterpart with catalytic residue conservation |
| P13798<br>461-732 | Q7Z5M8 | S587<br>D675<br>H707 | **S217 D304**<br>**H340** | 0.74 | UniProt: lipase activity<br>Pfam[53]: serine aminopeptidase domain ; structural similarity with prolyl oligopeptidase domain of reference counterpart with catalytic residue conservation |
| Q12884<br>503-760 | Q6P2H8 | S624<br>D702<br>H734 | **S113 D220**<br>**H252** | 0.71/ | UniProt: no MF reported<br>InterPro: has $\alpha/\beta$ hydrolase domain<br>Pfam: structural similarity with prolyl oligopeptidase domain of reference counterpart with catalytic residue conservation |
| **B. Active site residue predictions for novel human kinases against known references** |  |  |  |  |  |
| P11309 | Q92519 | D167,<br>K169 | **D175,**<br>**K177** | 0.67 | Has ability to bind and hydrolyze ATP weakly [54]<br>UniProt: annotation by similarity only; no experimental validation yet |
| Q8TCT0 | Q49MI3 | D197 | **D259** | 0.76 | UniProt: shows high structural similarity and catalytic conservation with known Ceramide kinase; no experimental work has been done to see if they bind to other kinase substrates except substrates that bind to ceramide kinases |
| Q00535 | Q59FN2 | D126 | **D126** | 0.74 | UniProt: non-specific serine/threonine kinase; annotation score 1/5; could be Cyclin-dependent kinase – based on high structural similarity |

**Table 2.** Model performance metrics for proteases and kinases model.

| **Table 2. Model performance metrics for proteases and kinases model.** |  |  |  |  |
| --- | --- | --- | --- | --- |
|  | Proteases |  | Kinases |  |
| Metrics | Pooled CV | Test | Pooled CV | Test |
| ROC-AUC | 0.98 | 0.99 | 0.96 | 0.98 |
| PR-AUC (avg. precision) | 0.98 | 0.97 | 0.97 | 0.99 |
| F1-score | 0.95 | 0.92 | 0.90 | 0.92 |
| precision | 0.98 | 0.90 | 0.90 | 1.00 |
| recall | 0.93 | 0.95 | 0.90 | 0.85 |
| Confusion matrix<br>(TN, FP, FN, TP) | (10479, 235,<br>745, 9469) | (31475, 295,<br>143, 2540) | (7556, 850,<br>898, 7789) | (1420, 0,<br>479, 2757) |

As summarized in **Table 1**, the identified reference-candidate pairs exhibit strong structural conservation, and also the identical amino acids of the catalytic pairs. Six representative pairs are illustrated in **Fig. 2** (for the other four, see **Supplementary Fig. S1**). Here, catalytic residues and their environment conservation are illustrated in **Fig.3**, where they show highly conserved local geometries.

**Figure 2.**
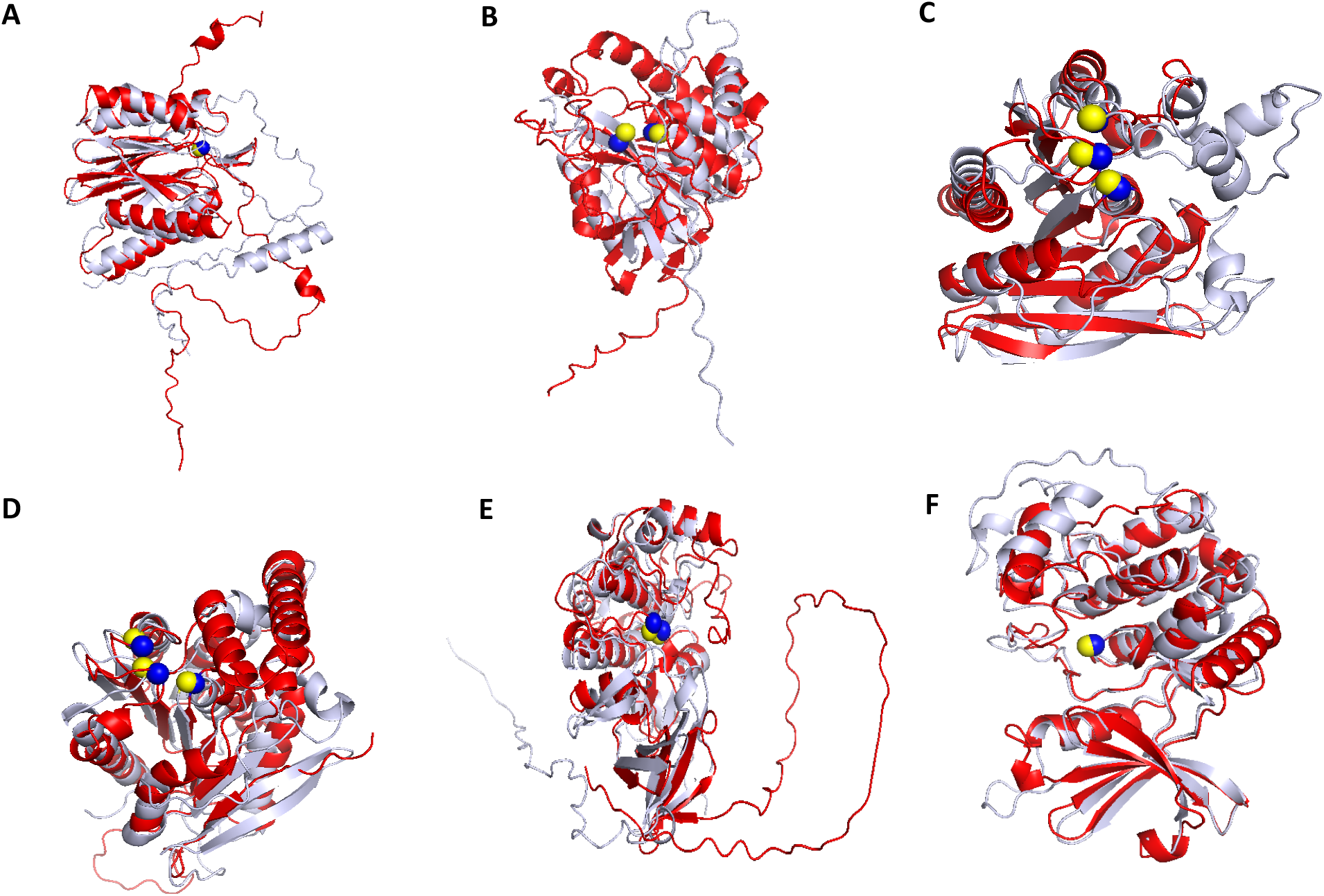
Pairwise superpositions of reference proteins with novel paralogs showing close positions of catalytic residues. Reference proteins are shown in gray and novel candidate proteins in red. Catalytic residues from reference and candidate proteins are shown as yellow and blue spheres, respectively. Catalytic residues for novel candidates are predicted through pairwise structural alignment using FoldMason[48] and visualizations with PyMOL. **(A-D)** Structural alignments of candidate proteases (**P28070, Q96NT3, Q5JUR7**, and **Q6P2H8)** with their corresponding reference proteins (**Table 1**) have RMSDs of 3.3 Å, 4.6 Å, 3.9 Å, and 4.2 Å, respectively. Here, proteins **Q5JUR7**, and **Q6P2H8** (**C** and **D**), both containing α/β hydrolase domains, are aligned with the prolyl oligopeptidase domain of the multi-domain reference proteins. **(E-F)** Candidate kinases **Q92519** and **Q59FN2** exhibit high structural similarity and catalytic residue conservation with their reference proteins (**Table 1**) with RMSDs of 3.0 Å and 2.6 Å. **Q92519** (Tribbles homolog 2, UniprotKB) is annotated to have MAP kinase regulatory activity, although earlier *in vitro* studies report weak kinase activity[54]. Similarly, **Q59FN2** (non-specific serine/threonine kinase protein, UniProtKB) is currently annotated as a kinase but is poorly characterized, lacking experimentally validated active-site annotations. Our independent analysis predicts both proteins to be candidate human kinases.

### More Precise Validation of Novel Paralogs

Finally, to further assess the functional plausibility of the predicted paralogs, we perform structural alignments on the catalytic residues between candidate proteins and their corresponding reference structures **(Fig. 3)**. Unlike conventional whole-protein superposition, which can obscure the structural metrics most relevant to function, these alignments on the catalytic residues make the most important comparison of functional geometric features. Despite structural divergence in distal regions, the candidate proteins show highly conserved spatial organization around catalytic residues with remarkably low active-site RMSDs, supporting preservation of mechanistically important features associated with the respective protease and kinase activities. Together, these results provide an additional layer of validation for the proposed novel paralogs.

**Figure 3.**
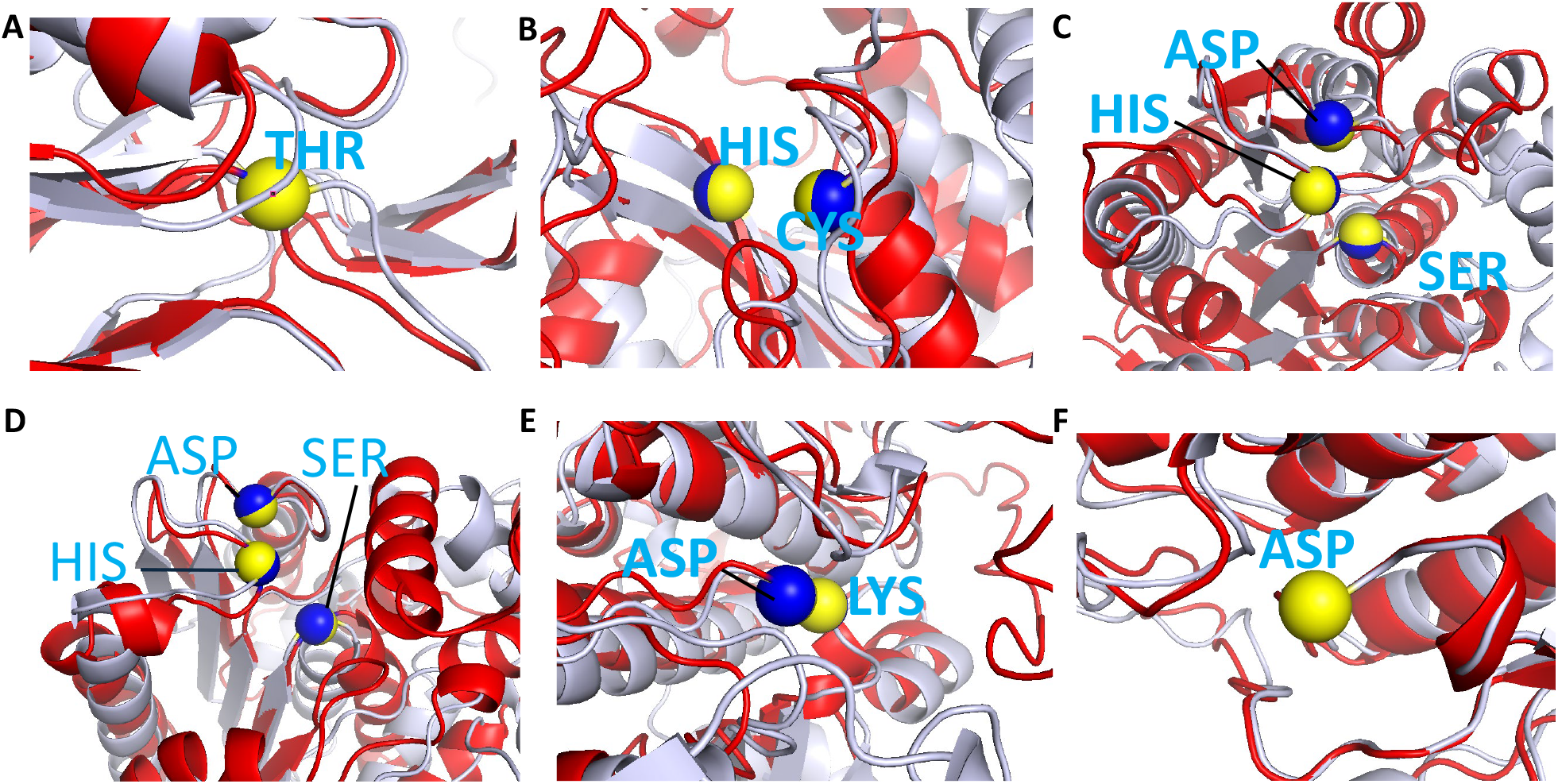
Structural alignments of novel candidates with reference proteins aligned on their identical catalytic C^α^ atoms. Active-site residues are colored as in **Fig. 2**. Some are not visible because they align nearly perfectly. This alignment sometimes increases deviations in other parts of the structures. **(A-D)** structural alignments of candidate proteases (P28070, Q96NT3, Q5JUR7, and Q6P2H8) with their reference proteases (**Table 1**), showing extremely close conserved catalytic residue alignments with RMSDs of 0.00 Å, 0.10 Å, 0.20 Å, and 0.06 Å respectively. **(E-F)** structural alignments of candidate kinases (Q92519, and Q59FN2) with their reference kinases (**Table 1**), exhibiting catalytic residue alignments with RMSDs of only 0.04 Å, and 0.00 Å respectively.

**Figure 4.**
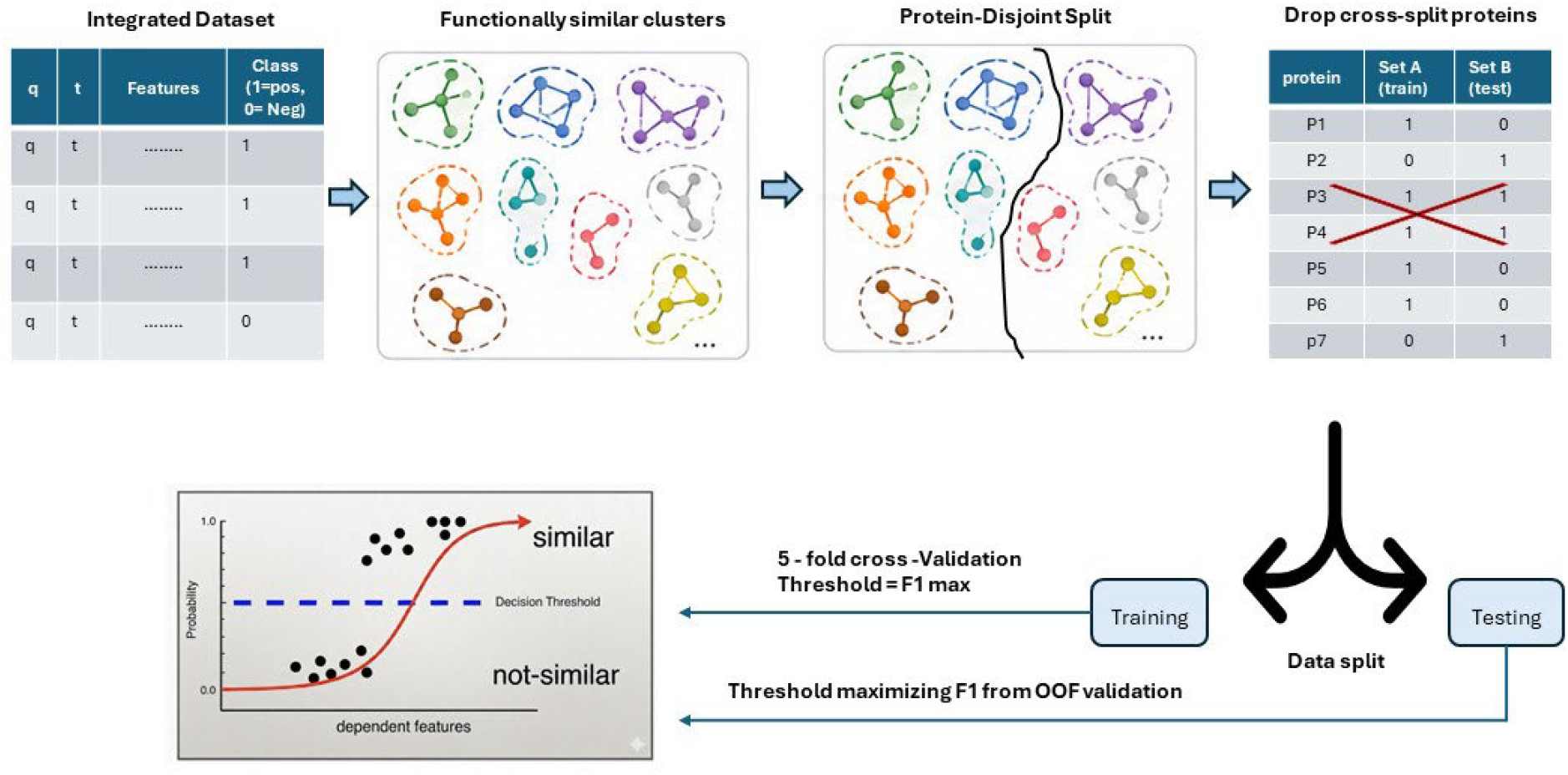
Protein data split and supervised logistic regression model training workflow. Protein pairs are represented as networks, where nodes correspond to individual proteins and edges are positive relationships, or functionally similar pairs. Connected components are partitioned into protein-disjoint training and test sets to prevent information leakage. Test datasets are comprised of 20% positive pairs and 40% from all proteins, while common members are excluded. Model training is performed using protein-disjoint paired 5-fold cross-validation. A threshold is chosen for each validation round that maximizes the F1-score. A final global threshold is determined from the pooled out-of-fold predictions and applied to the test dataset for model predictions. (See Supplementary material for additional details.)

**Figure 5.**
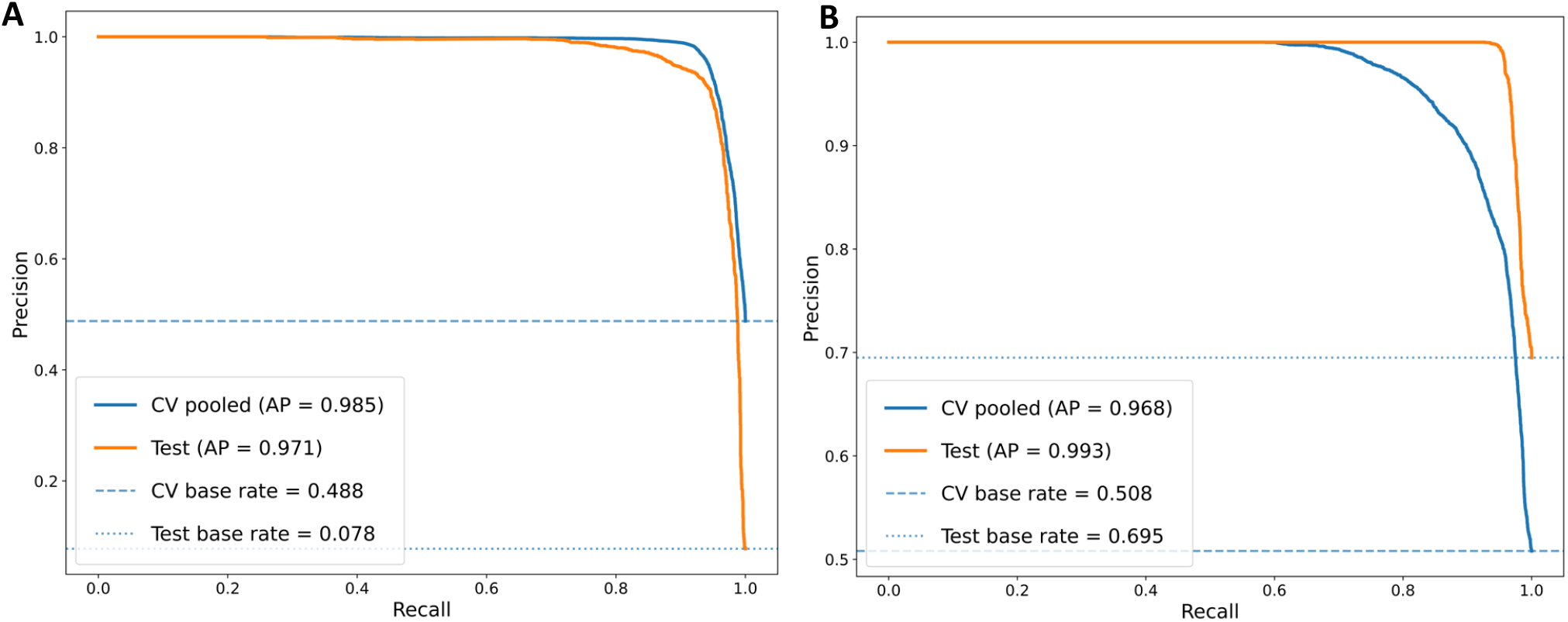
Model Performance Shown in Precision-Recall Curves: Proteases (A) and Kinases (B). Both perform well on clan-level and family-level characterization. The dashed horizontal lines represent the baseline precision corresponding to the class positive rate in the validation and test datasets highlighting the improved performance over random. Notably, the kinase model maintains higher precision across a wider recall range in the test dataset, suggesting kinase classification is comparatively easier because they are broadly conserved in sequence and structure at family level unlike the more stringent protease classifications.

## Conclusion and Discussion

Finding the novel protein functions at high resolution is fundamental for advancing protein annotations, accelerating drug discovery, and understanding molecular evolution. Traditionally, computational approaches for functional inference have relied on homolog searches based on sequence, structure, and more recently on protein language model-derived embedding similarities. In this study, we present a systematic framework for integrating complementary similarity scores from sequence – structure – embedding based homolog search methods to identify functionally related protein paralogs. By combining outputs from multiple homolog detection approaches, we develop two protein-family specific logistic regression models capable of distinguishing homolog pairs for functional relatedness. Using this approach together with downstream structural and catalytic residue conservation analyses, we predicted 7 novel protease paralogs and 3 novel kinase paralogs in humans. Notably, many of these proteins currently lack detailed functional annotations or experimentally validated active-site identifications for 8 of the 10 candidates in existing databases (see Table 1).

These results highlight an important limitation of conventional homolog-based annotation transfer: similarity alone is often insufficient to capture functional equivalence, particularly in protein families with diverse biochemical roles. For example, in the α/β hydrolases [40], serine carboxypeptidases and lipases differ functionally, but have closely similar structural folds and identical catalytic triads[41,42]. Here our findings demonstrate that combining family-specific predictions with regression modeling and detailed structural validation can substantially improve the identification of meaningful homolog relationships for more reliable function assignments.

Despite these promising results, several important challenges remain in supervised paralog identification and functional annotation transfer. First, supervised models rely heavily on the availability of high-quality annotations; however, many protein families and species remain poorly characterized without experimental or reliable computational validation, limiting construction of meaningful and reliable ground truth datasets. Second, functional similarity among proteins, particularly enzymes, should be most focused on the important residues, substrate-binding sites (and active sites for enzymes), and the local surrounding geometry, but such annotations remain incomplete or are entirely unavailable for many proteins. Third, predictive models are biased toward well-studied protein families, which confound annotation efforts. In contrast, under-studied proteins – arguably the most biologically important to annotate – are associated with sparse data, making both model training and downstream validation more difficult.

Importantly, humans represent one of the most extensively studied organisms, and both proteases and kinases are among the best-characterized enzyme families. Therefore, the relatively small number of novel candidates identified in this study could have been expected. Nevertheless, the protein family-specific approach developed here is particularly well-suited for application to less studied organisms, where it has the potential to uncover substantially larger numbers of homologs with functional identifications.

## Data and Methods

We obtain 675 human proteases from the MEROPS database and 522 kinases from the Kinhub database. After excluding kinases with ‘Atypical’ and ‘Other’ within Group names, we use 410 kinases. We choose protease functional labels from the MEROPS clan assignments and kinase group classifications from Kinhub and use these cases as the ground truth for supervised learning. To enable large-scale homolog discovery, we retrieve a total of 20,647 human protein sequences from the UniprotKB[14] reference proteome and the corresponding AlphaFold predicted structures (20,208 entries). Additionally, we collect the reviewed reference datasets of 536 proteases (EC 3.4) and 736 kinases (EC 2.7) from Swiss-Prot database to use for our novel paralog discovery and the downstream analysis.

### Integrated Homolog Search

We build method-specific databases by using BLASTp[16], MMseqs2, Foldseek[23], and PROST and perform all-vs-all exhaustive searches within each of the above datasets. BLASTp is an extremely well-known local sequence alignment-based homolog detection method, but surprisingly still yields some new homologs. MMseqs2[2] optimizes the alignment by first applying consecutive k-mer matches as a pre-filtering step, and it is 400 times faster and more sensitive than BLASTp. Foldseek does a structure-based homolog search by converting the three-dimensional structure information into a 1D alphabet string, followed by MMSeq2 sequence-based search on these strings that include structural information. PROST[5] is our own extremely different method that optimizes the use of the LPLM-based ESM1b [26] sequence embeddings to identify new homologs. Notably, PROST by itself performed particularly well in the CAFA 5 [35] competition for function prediction, scoring in the top 1.4 percentile. It performs well for remote homolog detection. In each dataset, we only keep the reciprocal hits from the homolog search and only the statistically significant matches are retained, using e-values less than 0.01. Finally, we prepare the integrated homolog dataset for each protein by combining the paralogs hit identified by each method.

### Imputing Missing Values and Dataset Augmentation

For each protein family, we generate all possible query-target pairs and extract similarity metrics from this integrated homolog pair dataset. These include sequence similarity metrics (percent identity, bitscore, e-value), structure similarity (TM-score), and embedding distance (PROST score). However, not all protein pairs are the hits from all methods. To deal with these missing values for the method-specific non-hits, we employ a controlled imputation strategy by assigning neutral or penalized values: for e-values and prost scores this is set equal to the threshold value; and for percent identity, bitscore, and TM-scores is set to zero. To mitigate the potential bias introduced by somewhat arbitrary values, we incorporate supplemental features (sequence_hit, structure_hit, prost_hit) to mask the method specific imputed values. Such features take binary values: 1 for real hits and 0 for non-hits. The target class or the truth value is defined as a binary value indicating whether the protein pair belongs to the same functional group, or not, i.e. MEROPS clan or kinase family.

### Protein-Disjoint Data Splitting, Training, and n-Fold Cross Validation

It is essential to divide the data into training, validation, and test sets in order to fit the model, tune and monitor performance, and to evaluate its generalization for unseen data. However, such evaluation is only meaningful when the model is stable, consistent, and free from overfitting. To prevent overfitting by information leakage, we devise a protein-disjoint data split based on graph connectivity. Proteins are represented as nodes in the graphs, and undirected edges connect proteins sharing the same functional label. As a result, connected components represent sets of proteins with functional coherence. To prevent leakage, entire clusters are assigned exclusively to either the training or the test set. However, a naïve random assignment of clusters can lead to class imbalance, as clusters vary widely in size and may disproportionally contribute positive (functionally related) pairs to one set. Additionally, because negative pairs are constructed from proteins within each split, the number of possible pairs decreases combinatorically with the number of proteins assigned to each set. To address these challenges, we adopt a controlled partitioning strategy: the two largest clusters are assigned to different splits, and the remaining clusters are distributed broadly to achieve an approximate 60:40 protein-level split between training and test sets, while maintaining a balanced 20% representation of positive pairs in the held-out test set.

The training set is further partitioned into training and validation subsets using the same protein-disjoint graph connectivity approach. To ensure robustness and reduce any dependence within any single functional cluster, we employ a protein-disjoint n-fold cross-validation scheme. In standard n-fold cross-validation, one-fold is held out for validation while the remaining folds are used for training. However, in our case, a single fold may contain proteins from only one cluster, resulting in validation sets not including any negative pairs. To overcome this, we use a paired-fold validation strategy, where two folds (corresponding to distinct clusters) are held out for validation, and the remaining n-2 folds are used for training. This ensures that both positive and negative pairs are present in each validation split.

During the model training, input features are scaled and class-weight balanced. We train the logistic regression model by minimizing the binary cross-entropy loss, *L*:

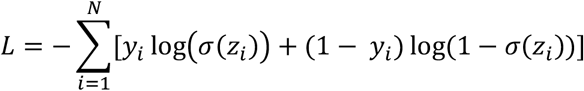

where *N* is the sample size (number of protein pairs), *y*_*i*_ *ϵ*{0, 1} is the true label indicating functional relatedness, and *z*_*i*_ = *W*^*T*^*X*_*scaled*_ + *b* is the linear model output, with *W* and *X* representing model weights and scaled feature vector for pair i, and b is the intercept. The σ function maps the linear output to the probability score between 0 and 1.

For each cross-validation round, an optimal threshold is selected by maximizing the F1-score on the corresponding validation set. Subsequently, we pool all the predictions from validation folds to form an out-of-fold (OOF) dataset, from which a global threshold is determined by maximizing the overall F1-score. This threshold is then applied to the held-out protein-disjoint test set for model evaluation.

### Model Evaluation

Finally, the trained model is used for the held-out test dataset to measure how well it performs on untrained data. We show the model performance values for the pooled validation and test sets in terms of the various metrics, precision, recall, F1-score, average precision (PR AUC), and ROC AUC as shown in the table below. Detailed values as well as model data splits are shown in **Supplementary Table S1 and S2**.

## Authors’ contribution

**Pradeep BK:** Writing – original draft, Writing – review & editing. **Weixia Deng:** Supported data analysis. Writing – review. **Robert L. Jernigan:** Supervision, Funding acquisition, Writing – review & editing. All authors read and approved the final manuscript.

## Supplementary material

### Data and Code Availability

Data and code are available at https://github.com/pradeepbkarma/paralog-discovery-pipeline

## Acknowledgements

This work was supported by NIH grants R01HG012117, R01GM144961, and R01GM157600.

## References

[1] Amaral P, Carbonell-Sala S, De La Vega FM, Faial T, Frankish A, Gingeras T, et al. The status of the human gene catalogue. Nature 2023;622:41–7. 10.1038/S41586-023-06490-X.

[2] Steinegger M, Söding J. MMseqs2 enables sensitive protein sequence searching for the analysis of massive data sets. Nat Biotechnol 2017;35:1026–8. 10.1038/NBT.3988;SUBJMETA=114,171,2184,2404,326,631,794;KWRD=ENVIRONMENTAL+MICROBIOLOGY,FUNCTIONAL+CLUSTERING,SEQUENCE+ANNOTATION,SOFTWARE.

[3] Baltoumas FA, Karatzas E, Paez-Espino D, Venetsianou NK, Aplakidou E, Oulas A, et al. Exploring microbial functional biodiversity at the protein family level—From metagenomic sequence reads to annotated protein clusters. Frontiers in Bioinformatics 2023;3:1157956. 10.3389/FBINF.2023.1157956/FULL.

[4] Gan HH, Perlow RA, Roy S, Ko J, Wu M, Huang J, et al. Analysis of protein sequence/structure similarity relationships. Biophys J 2002;83:2781–91. 10.1016/s0006-3495(02)75287-9.

[5] Kilinc M, Jia K, Jernigan RL. Improved global protein homolog detection with major gains in function identification. Proc Natl Acad Sci U S A 2023;120:e2211823120. 10.1073/PNAS.2211823120/SUPPL_FILE/PNAS.2211823120.SD01.XLSX.

[6] Jumper J, Evans R, Pritzel A, Green T, Figurnov M, Ronneberger O, et al. Highly accurate protein structure prediction with AlphaFold. Nature 2021;596:583–9. 10.1038/s41586-021-03819-2.

[7] Jumper J, Hassabis D. Protein structure predictions to atomic accuracy with AlphaFold. Nature Methods 2022 19:1 2022;19:11–2. 10.1038/s41592-021-01362-6.

[8] Lin Z, Akin H, Rao R, Hie B, Zhu Z, Lu W, et al. Evolutionary-scale prediction of atomic-level protein structure with a language model. Science (1979) 2023;379:1123–30. 10.1126/SCIENCE.ADE2574/SUPPL_FILE/SCIENCE.ADE2574_SM.PDF.

[9] Baek M, DiMaio F, Anishchenko I, Dauparas J, Ovchinnikov S, Lee GR, et al. Accurate prediction of protein structures and interactions using a three-track neural network. Science (1979) 2021;373:871–6. 10.1126/SCIENCE.ABJ8754/SUPPL_FILE/ABJ8754_MDAR_REPRODUCIBILITY_CHECKLIST.PDF.

[10] Pearce R, Li Y, Omenn GS, Zhang Y. Fast and accurate Ab Initio Protein structure prediction using deep learning potentials. PLoS Comput Biol 2022;18:e1010539. 10.1371/JOURNAL.PCBI.1010539.

[11] Wu R, Ding F, Wang R, Shen R, Zhang X, Luo S, et al. High-resolution de novo structure prediction from primary sequence. BioRxiv 2022:2022.07.21.500999. 10.1101/2022.07.21.500999.

[12] Brandes N, Ofer D, Peleg Y, Rappoport N, Linial M. ProteinBERT: a universal deep-learning model of protein sequence and function. Bioinformatics 2022;38:2102–10. 10.1093/BIOINFORMATICS/BTAC020.

[13] De Crécy-Lagard V, Amorin De Hegedus R, Arighi C, Babor J, Bateman A, Blaby I, et al. A roadmap for the functional annotation of protein families: a community perspective. Database 2022;2022. 10.1093/DATABASE/BAAC062.

[14] Bateman A, Martin MJ, Orchard S, Magrane M, Adesina A, Ahmad S, et al. UniProt: the Universal Protein Knowledgebase in 2025. Nucleic Acids Res 2025;53:D609–17. 10.1093/NAR/GKAE1010.

[15] Bairoch A, Apweiler R. The SWISS-PROT protein sequence database and its supplement TrEMBL in 2000. Nucleic Acids Res 2000;28:45. 10.1093/NAR/28.1.45.

[16] Altschul SF, Gish W, Miller W, Myers EW, Lipman DJ. Basic local alignment search tool. J Mol Biol 1990;215:403–10. 10.1016/S0022-2836(05)80360-2.

[17] Edgar RC. Search and clustering orders of magnitude faster than BLAST. Bioinformatics 2010;26:2460–1. 10.1093/BIOINFORMATICS/BTQ461.

[18] Zhao Y, Tang H, Ye Y. RAPSearch2: a fast and memory-efficient protein similarity search tool for next-generation sequencing data. Bioinformatics 2012;28:125–6. 10.1093/BIOINFORMATICS/BTR595.

[19] Buchfink B, Xie C, Huson DH. Fast and sensitive protein alignment using DIAMOND. Nature Methods 2014 12:1 2014;12:59–60. 10.1038/nmeth.3176.

[20] Holm L, Kääriäinen S, Rosenström P, Schenkel A. Searching protein structure databases with DaliLite v.3. Bioinformatics 2008;24:2780–1. 10.1093/BIOINFORMATICS/BTN507.

[21] Zhang Y, Skolnick J. TM-align: A protein structure alignment algorithm based on the TM-score. Nucleic Acids Res 2005;33:2302–9. 10.1093/NAR/GKI524,.

[22] Shindyalov IN, Bourne PE. Protein structure alignment by incremental combinatorial extension (CE) of the optimal path. Protein Eng 1998;11:739–47. 10.1093/PROTEIN/11.9.739.

[23] van Kempen M, Kim SS, Tumescheit C, Mirdita M, Lee J, Gilchrist CLM, et al. Fast and accurate protein structure search with Foldseek. Nat Biotechnol 2024;42:243–6. 10.1038/S41587-023-01773-0;SUBJMETA=114,535,631,794;KWRD=COMPUTATIONAL+BIOLOGY+AND+BIOINFORMATICS,SOFTWARE,STRUCTURAL+BIOLOGY.

[24] Knudsen M, Wiuf C. The CATH database. Hum Genomics 2010;4:207. 10.1186/1479-7364-4-3-207.

[25] Kilinc M, Jia K, Jernigan RL. Major advances in protein function assignment by remote homolog detection with protein language models – A review. Curr Opin Struct Biol 2025;90:102984. 10.1016/J.SBI.2025.102984.

[26] Rives A, Meier J, Sercu T, Goyal S, Lin Z, Liu J, et al. Biological structure and function emerge from scaling unsupervised learning to 250 million protein sequences. Proc Natl Acad Sci U S A 2021;118:e2016239118. 10.1073/PNAS.2016239118/SUPPL_FILE/PNAS.2016239118.SAPP.PDF.

[27] Hayes T, Rao R, Akin H, Sofroniew NJ, Oktay D, Lin Z, et al. Simulating 500 million years of evolution with a language model. Science (1979) 2025;387:850–8. 10.1126/SCIENCE.ADS0018;ISSUE:ISSUE:DOI.

[28] Elnaggar A, Heinzinger M, Dallago C, Rehawi G, Wang Y, Jones L, et al. ProtTrans: Toward Understanding the Language of Life Through Self-Supervised Learning. IEEE Trans Pattern Anal Mach Intell 2022;44:7112–27. 10.1109/TPAMI.2021.3095381.

[29] Brandes N, Ofer D, Peleg Y, Rappoport N, Linial M. ProteinBERT: a universal deep-learning model of protein sequence and function. Bioinformatics 2022;38:2102–10. 10.1093/BIOINFORMATICS/BTAC020.

[30] Madani A, Mccann B, Naik N, Shirish Keskar N, Anand N, Chu A, et al. ProGen: Language Modeling for Protein Generation n.d.

[31] Liu W, Wang Z, You R, Xie C, Wei H, Xiong Y, et al. PLMSearch: Protein language model powers accurate and fast sequence search for remote homology. Nature Communications 2024 15:1 2024;15:2775–. 10.1038/s41467-024-46808-5.

[32] Keluskar A, Batra P, Bezshapkin V, Morton JT, Zhu Q. TM-Vec 2: Accelerated Protein Homology Detection for Structural Similarity. BioRxiv 2026:2026.02.05.704073. 10.64898/2026.02.05.704073.

[33] Heinzinger M, Weisseno w K, Sanchez JG, Henkel A, Mirdita M, Steinegger M, et al. Bilingual language model for protein sequence and structure. NAR Genom Bioinform 2024;6:150. 10.1093/NARGAB/LQAE150.

[34] Xiao Y, Sun E, Jin Y, Wang Q, Wang W. ProteinGPT: Multimodal LLM for Protein Property Prediction and Structure Understanding 2024.

[35] Kaluza MCDP, Ramola R, Joshi P, Piovesan D, Reade W, Orchard S, et al. Advances in Protein Function Prediction from the Fifth CAFA Challenge. BioRxiv 2026:2026.04.27.716980. 10.64898/2026.04.27.716980.

[36] Leclercq M, Droit A. Protein Language Models: Applications and Perspectives. J Proteome Res 2025;25:507. 10.1021/ACS.JPROTEOME.5C00506.

[37] Marques AC, Vinckenbosch N, Brawand D, Kaessmann H. Functional diversification of duplicate genes through subcellular adaptation of encoded proteins. Genome Biol 2008;9:R54. 10.1186/GB-2008-9-3-R54.

[38] Liu ZP, Wu LY, Wang Y, Zhang XS, Chen L. Bridging protein local structures and protein functions. Amino Acids 2008;35:627. 10.1007/S00726-008-0088-8.

[39] Cagiada M, Bottaro S, Lindemose S, Schenstrøm SM, Stein A, Hartmann-Petersen R, et al. Discovering functionally important sites in proteins. Nature Communications 2023 14:1 2023;14:4175–. 10.1038/s41467-023-39909-0.

[40] Ozhelvaci F, Steczkiewicz K. α/β Hydrolases: Toward Unraveling Entangled Classification. Proteins: Structure, Function and Bioinformatics 2025;93:855–70. 10.1002/PROT.26776;WGROUP:STRING:PUBLICATION.

[41] Miled N, Bussetta C, De Caro A, Rivière M, Berti L, Canaan S. Importance of the lid and cap domains for the catalytic activity of gastric lipases. Comp Biochem Physiol B Biochem Mol Biol 2003;136:131–8. 10.1016/S1096-4959(03)00183-0.

[42] Roussel A, Canaan S, Egloff MP, Rivière M, Dupuis L, Verger R, et al. Crystal structure of human gastric lipase and model of lysosomal acid lipase, two lipolytic enzymes of medical interest. Journal of Biological Chemistry 1999;274:16995–7002. 10.1074/jbc.274.24.16995.

[43] Yan H, Wang S, Liu H, Mamitsuka H, Zhu S. GORetriever: reranking protein-description-based GO candidates by literature-driven deep information retrieval for protein function annotation. Bioinformatics 2024;40:ii53–61. 10.1093/BIOINFORMATICS/BTAE401.

[44] Senoner T, Vahidi P, Olenyi T, Senoner F, Sisman G, Kahl E, et al. ProtSpace: Protein Universe in Your Browser. BioRxiv 2026:2026.05.04.722720. 10.64898/2026.05.04.722720.

[45] Sperandei S. Understanding logistic regression analysis. Biochem Med (Zagreb) 2014;24:12. 10.11613/BM.2014.003.

[46] Rawlings ND, Tolle DP, Barrett AJ. MEROPS: the peptidase database. Nucleic Acids Res 2004;32:D160. 10.1093/NAR/GKH071.

[47] Eid S, Turk S, Volkamer A, Rippmann F, Fulle S. KinMap: a web-based tool for interactive navigation through human kinome data. BMC Bioinformatics 2017 18:1 2017;18:16–. 10.1186/S12859-016-1433-7.

[48] Gilchrist CLM, Mirdita M, Steinegger M. Multiple protein structure alignment at scale with FoldMason. Science 2026;391:485–8. 10.1126/SCIENCE.ADS6733.

[49] Becker AB, Roth RA. An unusual active site identified in a family of zinc metalloendopeptidases. Proc Natl Acad Sci U S A 1992;89:3835–9. 10.1073/PNAS.89.9.3835.

[50] Taylor AB, Smith BS, Kitada S, Kojima K, Miyaura H, Otwinowski Z, et al. Crystal structures of mitochondrial processing peptidase reveal the mode for specific cleavage of import signal sequences. Structure 2001;9:615–25. 10.1016/S0969-2126(01)00621-9.

[51] Bode W, Gomis-Rüth FX, Stöckler W. Astacins, serralysins, snake venom and matrix metalloproteinases exhibit identical zinc-binding environments (HEXXHXXGXXH and Met-turn) and topologies and should be grouped into a common family, the “metzincins.” FEBS Lett 1993;331:134–40. 10.1016/0014-5793(93)80312-I.

[52] Blum M, Andreeva A, Florentino LC, Chuguransky SR, Grego T, Hobbs E, et al. InterPro: the protein sequence classification resource in 2025. Nucleic Acids Res 2025;53:D444–56. 10.1093/NAR/GKAE1082.

[53] Mistry J, Chuguransky S, Williams L, Qureshi M, Salazar GA, Sonnhammer ELL, et al. Pfam: The protein families database in 2021. Nucleic Acids Res 2021;49:D412–9. 10.1093/NAR/GKAA913.

[54] Bailey FP, Byrne DP, Oruganty K, Eyers CE, Novotny CJ, Shokat KM, et al. The Tribbles 2 (TRB2) pseudokinase binds to ATP and autophosphorylates in a metal-independent manner. Biochem J 2015;467:47–62. 10.1042/BJ20141441.

