## Supplemental Information for "New Protein Function Characterization for Human Paralog Discovery, Scraping the Bottom of the Genomics Barrel"

Supplementary Information

Data- split

| **Table S1. Data split in training and testing set for MEROPS and Kinhub logistic regression model.** | | |
|---|---|---|
| | Proteases – MEROPS | Kinases – Kinhub |
| Total proteins | 675 | 398 |
| Proteins in training set | 412 | 301 |
| Proteins in test set | 263 | 97 |
| Training rows | 84666 | 45150 |
| Test rows | 34453 | 4656 |
| Train positives | 10214 | 9585 |
| Test positives | 2683 | 3236 |

model comparison summary

| **Table 2. Model performance metrices for protease and kinase family specific regression model.** | | | | |
|---|---|---|---|---|
| | Proteases | | Kinases | |
| | Validation | Test | Validation | Test |
| Accuracy | 0.95 | 0.98 | 0.89 | 0.90 |
| Precision | 0.97 | 0.89 | 0.90 | 1.00 |
| Recall | 0.93 | 0.95 | 0.90 | 0.85 |
| F1 | 0.95 | 0.92 | 0.90 | 0.92 |
| Average precision | 0.98 | 0.97 | 0.97 | 0.99 |
| ROC AUC | 0.98 | 0.99 | 0.96 | 0.98 |
| TN | 10479 | 31475 | 7556 | 1420 |
| FP | 235 | 295 | 850 | 0 |
| FN | 745 | 143 | 898 | 479 |
| TP | 9469 | 2540 | 7784 | 2757 |

$$Precision = \frac{TP}{TP + FP}$$

$$Recall = \frac{TP}{TP + FN}$$

$$F1\ score = 2\ \frac{Precision\ \times Recall}{Precision + Recall}$$

$$AUC = \int_0^1 TPR(FPR)dFPR$$

Here, TP = True positive, FP = False positive, FN = False negative, TN = True Negative, AUC is Area under the curve formed by plotting False Positive Rate (FPR) and True Positive Rate (TPR), to approximate the definite integral. AUC-ROC is used to check how well the classifier discriminates the positive cases from negative cases.

$$\text{TM-score} = Max\left[\frac{1}{L_{\text{Target}}}\sum_{i}^{L_{\text{ali}}} \frac{1}{1+\left(\frac{d_i}{d_0(L_{\text{Target}})}\right)^2}\right]$$

Here, di is the distance between the aligned residues, $d_0(L_{\text{Target}}) = 1.24\sqrt[3]{L_{Target} - 15} - 1.8$ is a distance parameter - normalizes the distance so that the average TM-score is not dependent on the protein size for random structure pairs, $L_{ali}$ is the length of aligned residues.

Figure SA. P27487 Res(497-766) in bluewhite and Q7L211 in red

**A**

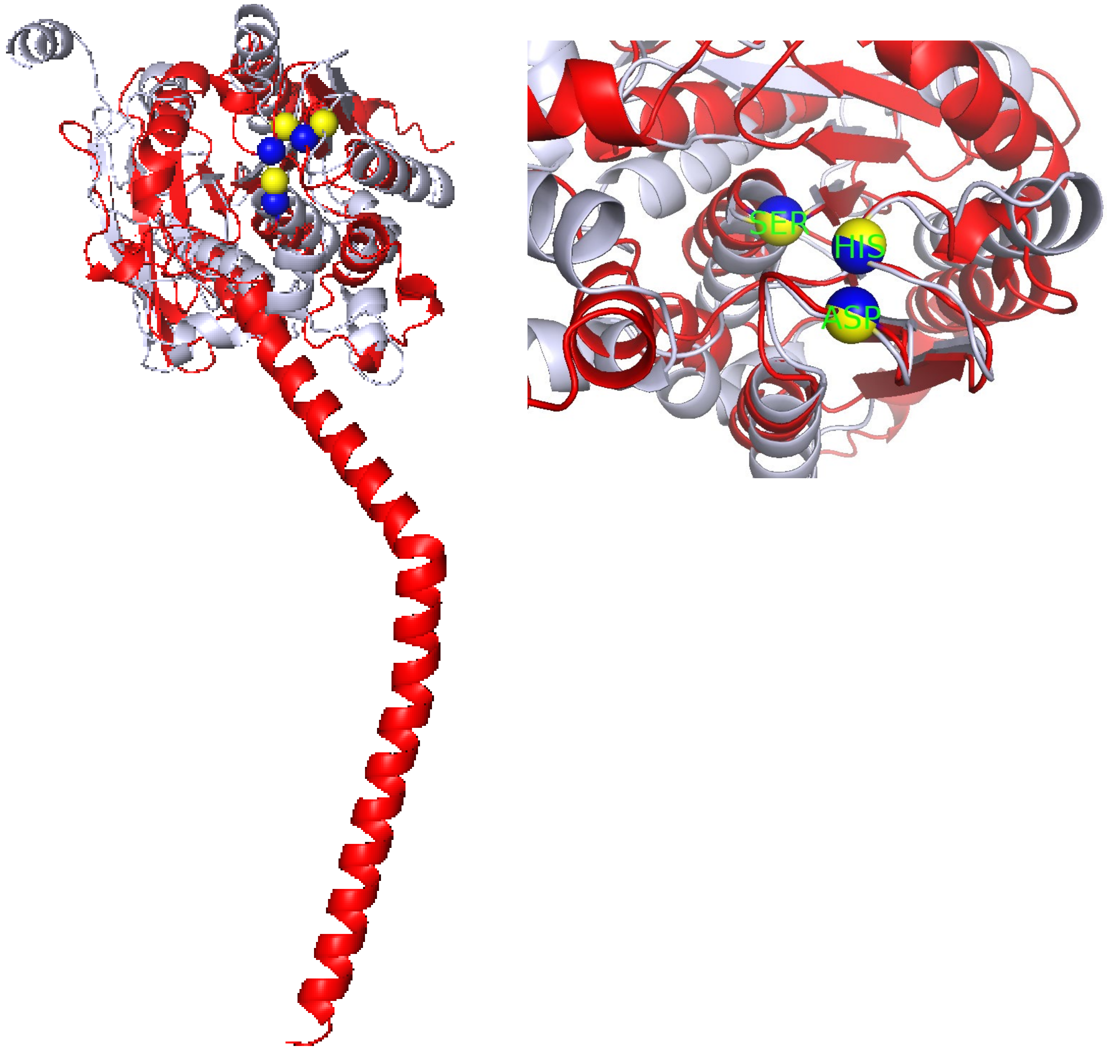

**B**

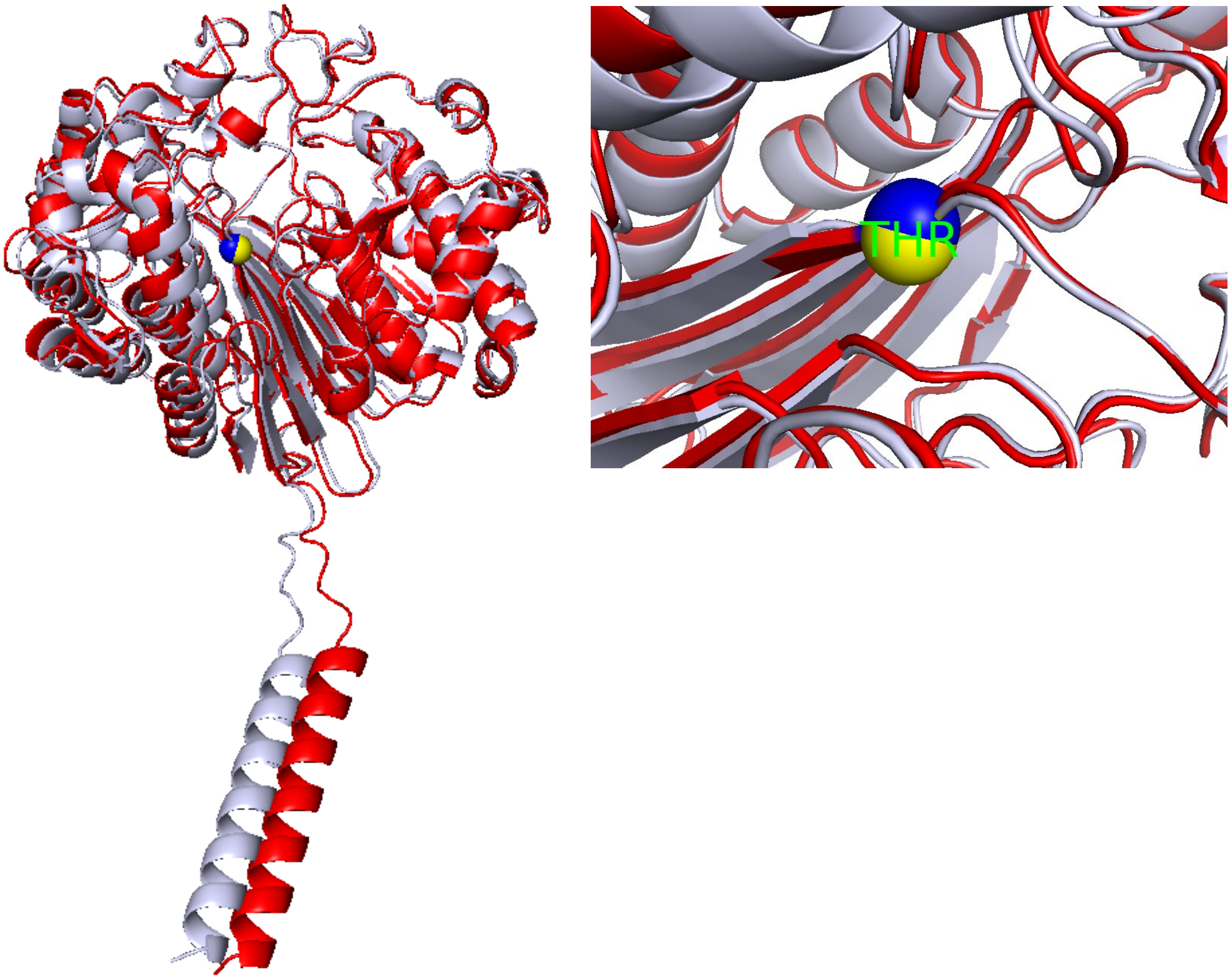


**C**

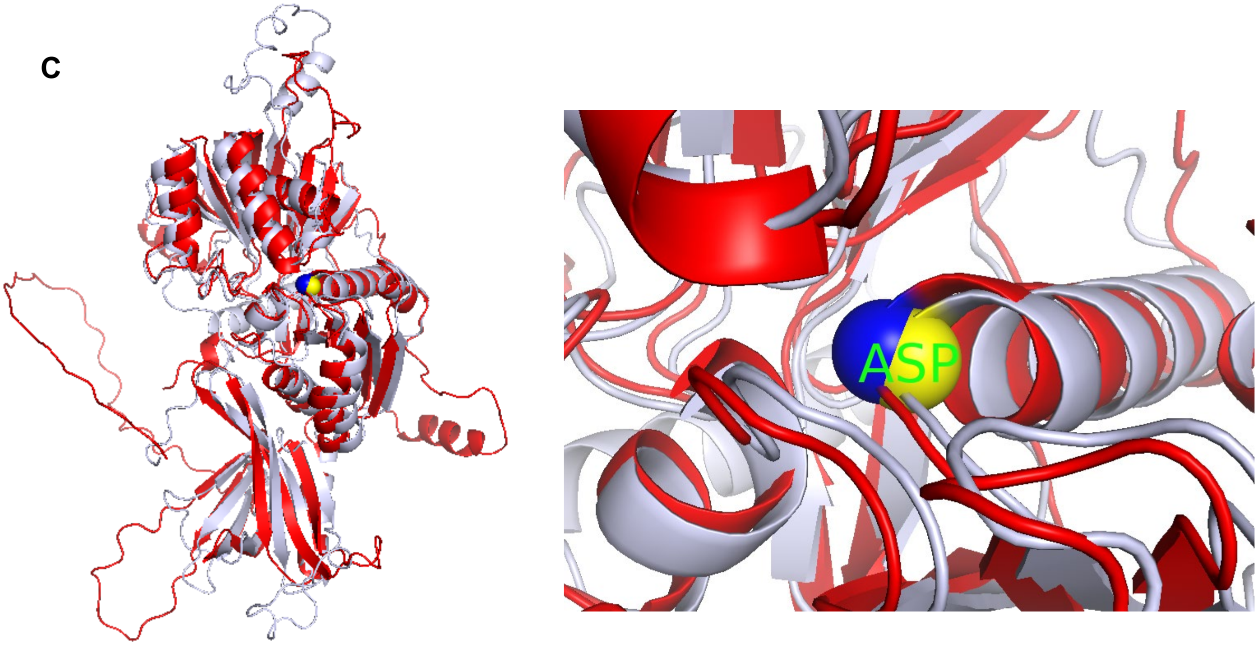

D

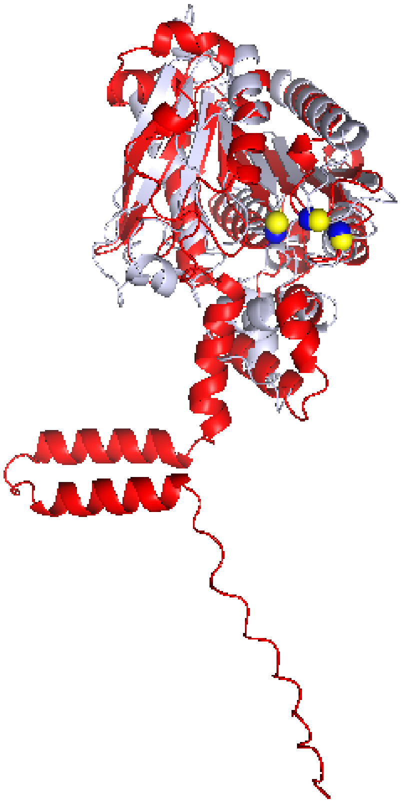

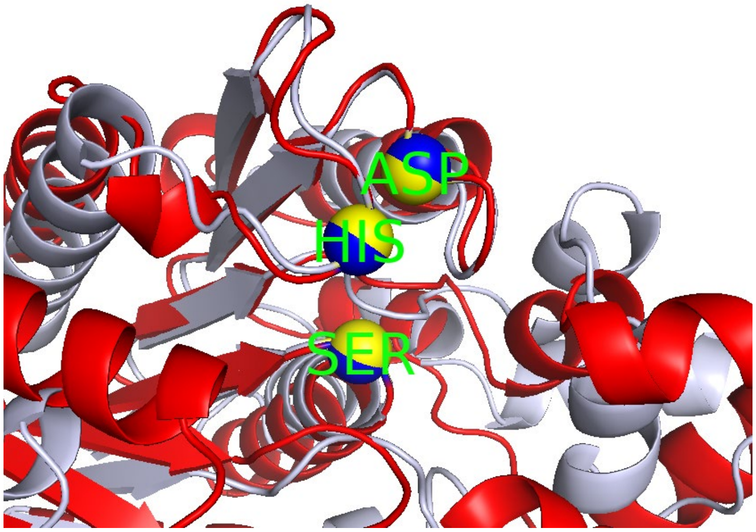


Figure S1. **Pairwise structural superposition of reference proteins and novel paralog candidates reveals conservation of catalytic residues.** Reference proteins are shown in blue-white and novel candidate proteins in red. Catalytic residues from reference and candidate proteins are shown in yellow and blue spheres, respectively. Catalytic residues for putative novel candidates are predicted through pairwise structural alignment using FoldMason and structural visualizations are generated using PyMOL**.** **A.** Novel protease candidate **Q7L211** aligned with reference protease P27487 Res(497-766) with RMSD of 4.6 Å, and subsequent alignment on their catalytic triad with RMSD of 0.12 Å. **B**. Novel protease candidate **P36268** aligned with corresponding reference protein P19440 with RMSD of 1.52 Å when aligned globally. **C**. Putative kinase protein **Q49MI3** aligned with known kinase reference protein Q8TCT0 with RMSD of 4.66 Å. **D.** Novel protease candidate Q7Z5M8 aligned with reference protease P13798 with RMSD of 5.5 Å and subsequent alignment on their catalytic triad with RMSD of 0.13 Å.

Model weights

| feature | coef | abs_coef | odds_ratio_per_1sd | direction |
| --- | --- | --- | --- | --- |
| bitscore | 5.9015 | 5.9015 | 365.5698 | increases P(class=1) |
| tm_gm | 2.6293 | 2.6293 | 13.8646 | increases P(class=1) |
| p_score | -0.6945 | 0.6945 | 0.4993 | decreases P(class=1) |
| evalue | -0.5204 | 0.5204 | 0.5943 | decreases P(class=1) |
| q_tm | 0.4589 | 0.4589 | 1.5823 | increases P(class=1) |
| seq_hit | -0.1489 | 0.1489 | 0.8616 | decreases P(class=1) |
| pident | -0.034 | 0.034 | 0.9666 | decreases P(class=1) |
| str_hit | 0.0 | 0.0 | 1.0 | increases P(class=1) |
| prost_hit | 0.0 | 0.0 | 1.0 | increases P(class=1) |
